# Activation of Cytochrome C Peroxidase Function Through Coordinated Foldon Loop Dynamics upon Interaction with Anionic Lipids

**DOI:** 10.1101/2021.02.24.432556

**Authors:** Mingyue Li, Wanyang Sun, Vladimir A. Tyurin, Maria DeLucia, Jinwoo Ahn, Valerian E. Kagan, Patrick C.A. van der Wel

## Abstract

Cardiolipin (CL) is a mitochondrial anionic lipid that plays important roles in the regulation and signaling of mitochondrial apoptosis. CL peroxidation catalyzed by the assembly of CL-cytochrome c (cyt c) complexes at the inner mitochondrial membrane is a critical checkpoint. The structural changes in the protein, associated with peroxidase activation by CL and different anionic lipids, are not known at a molecular level. To better understand these peripheral protein-lipid interactions, we compare how phosphatidylglycerol (PG) and CL lipids trigger cyt c peroxidase activation, and correlate functional differences to structural and motional changes in membrane-associated cyt c. Structural and motional studies of the bound protein are enabled by magic angle spinning solid state NMR spectroscopy, while lipid peroxidase activity is assayed by mass spectrometry. PG binding results in a surface-bound state that preserves a nativelike fold, which nonetheless allows for significant peroxidase activity, though at a lower level than binding its native substrate CL. Lipid-specific differences in peroxidase activation are found to correlate to corresponding differences in lipid-induced protein mobility, affecting specific protein segments. The dynamics of omega loops C and D are upregulated by CL binding, in a way that is remarkably controlled by the protein:lipid stoichiometry. In contrast to complete chemical denaturation, membrane-induced protein destabilization reflects a destabilization of select cyt c foldons, while the energetically most stable helices are preserved. Our studies illuminate the interplay of protein and lipid dynamics in the creation of lipid peroxidase-active proteolipid complexes implicated in early stages of mitochondrial apoptosis.

## INTRODUCTION

Cytochrome c (cyt c) is a multifunctional protein whose primary role is to shuttle electrons in the respiratory chain within the intermembrane space (IMS) of mitochondria [1]. Cyt c is also increasingly known for its gain of function as a lipid peroxidase upon interaction with the mitochondria-specific phospholipid cardiolipin (CL) in regulating intrinsic apoptosis [2]. Intrinsic apoptosis is a highly regulated cell death pathway in which mitochondria play an essential role in the elimination of unwanted or unhealthy cells, in response to internal stimuli such as DNA damage, metabolic stress, and unfolded proteins [3]. Pathogenic dysregulation of apoptosis is associated with tumorigenesis as well as neurodegenerative diseases, such as Alzheimer’s and Huntington’s disease [4, 5]. Early steps of intrinsic apoptosis involve a redistribution and peroxidation of CL in mitochondrial membranes [3, 6]. In normal cells, CL is almost exclusively found in the mitochondrial inner membrane (MIM) where it fulfills crucial roles in maintaining mitochondrial function and MIM organization. Under apoptotic conditions, the regulated distribution of CL across the inner and outer mitochondrial membranes is disrupted. Among other effects, this makes an increasing amount of CL available for binding cyt c at the IMS membrane surface. Tight binding between the positively charged cyt c and these negatively charged lipids creates a protein-lipid nanocomplex with distinct properties from native cyt c [7]. Crucially, this proteolipid complex catalyzes the specific oxidation of polyunsaturated CL by reactive oxygen species (ROS) [2, 8]. The targeted lipid peroxidation is then a precursor to mitochondrial outer membrane permeabilization (MOMP) and cyt c release.

Structurally, horse heart cyt c contains 5 helices (named the N, 50’s, 60’s, 70’s, and C helices) spaced by Ω loops B, C, D from the N to C terminus (**Figure 1a**). The functional state of the protein contains a covalently attached heme, which is partially encapsulated and stabilized by axial ligands Met80 of Ω loop D and His18 of Ω loop B (**Figure 1b**). Beyond its essential role in many cellular processes, historical studies of cyt c have also revealed important biophysical principles governing protein folding and function [1, 9]. Native-state hydrogen exchange experiments elucidated the folding pathway of cyt c, leading to the definition of the five cooperative folding units, known as foldons, that form the tertiary structure [10–13]. These foldons undergo cooperative folding along a stepwise, energetically favorable pathway. The different foldons are often identified by the color codes represented in **Figure 1a** and **1b** [10]. Destabilization or modulation of this native fold causes a change in the cyt c functional properties, explaining at least in part how CL binding induces a boost in peroxidase activity during apoptosis [1, 14]. Yet, exactly how the fold and dynamics of cyt c are modified upon membrane binding has remained difficult to assess, given the apparent heterogeneity and dynamics of the lipid-bound protein and the technical challenges of probing peripherally bound proteins in atomic detail.

**Figure 1.**
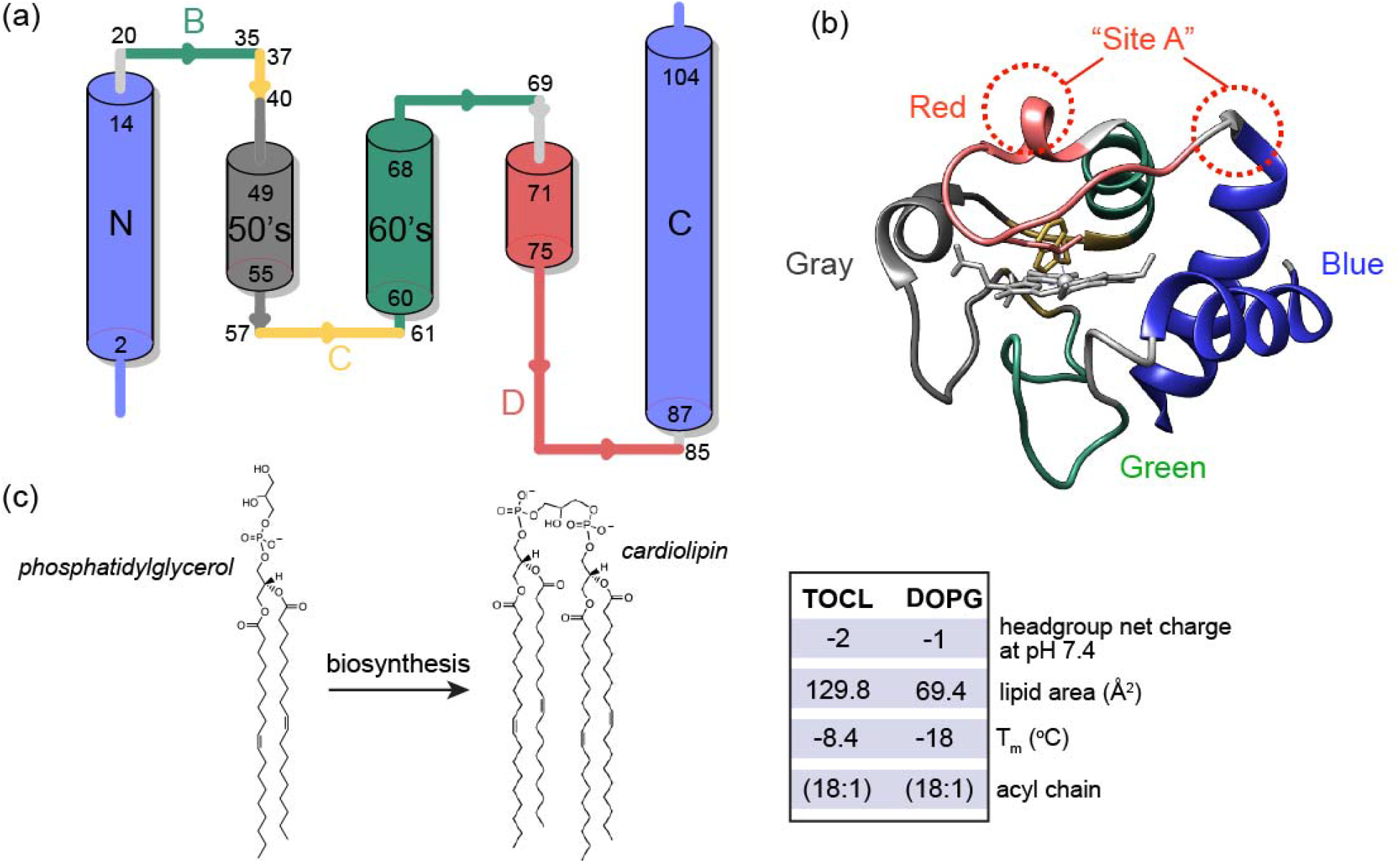
Cytochrome c and its anionic lipid partners. (a) Cartoon representation of cyt c secondary structure, showing the five foldons [34]: the blue foldon (N and C-terminal helices, residues 1-14 and 88-104), the green foldon (helix 61-69 and Ω loop 20-35), the gray foldon (Ω loop 40-57), the red foldon (Ω loop 71-85) and the yellow unit (residues 37-40 and 57-60). (b) 3D structure of horse heart cyt c [35] (PDB ID: 1hrc) colored and labeled according to the different foldons [10]. The dotted red circle marks the “site A” lipid binding site. (c) Chemical structure and enzymatic interconversion of phosphatidylglyerol (PG) and CL, catalyzed by CL synthase. (d) Comparison of headgroup net charge, lipid area, liquid to gel phase transition temperature (T_m_) and the acyl chain information of TOCL and DOPG [36–38].

The primary native substrates of cyt c’s pro-apoptotic peroxidase activity are poly-unsaturated CL species [2]. CL represents a unique class of anionic mitochondrial phospholipids, with essential functions in mitochondrial respiration, remodeling, mitophagy, and apoptosis [15–17]. A reduced level of CL in mitochondria usually leads to the disruption of ATP synthesis and loss of cristae morphology [18]. Disruption of CL biosynthesis and metabolism is pathogenic, as exemplified in Barth syndrome [19]. The biosynthesis of CL shares similar pathways with other phospholipids as it passes through common intermediates. Only the last step of CL synthesis is a unique reaction, in which cardiolipin synthase catalyzes the conversion of phosphatidylglycerol (PG) to CL (**Figure 1c**) [16, 20]. In the case of CL abnormalities, some lost CL functions can be rescued in part by this PG precursor, indicative of its similarities in chemical structure, (anionic) charge and (small) headgroup size (**Figure 1d**) [18, 21–25]. A yeast mutant which lacks CL synthase and has no detectable CL, accumulates PG and displays normal mitochondrial activity. PG incorporates into mitochondrial membranes to reduce mitochondrial dysfunction due to CL deficiency [21]. It is proposed that PG can act as a substitute for CL for its essential functions in the respiratory chain, except when cells are under stress [22, 26]. *In vitro*, there are many studies reporting on the interactions of cyt c with non-CL anionic lipids, including PG, showing the potential for similar interactions to those underpinning CL-induced peroxidase activation [27–30]. Yet, to the best of our knowledge there is no record of a direct involvement of PG in place of CL in the apoptotic pathway *in vivo*, with studies showing CL to be the preferred substrate for cyt c binding and peroxidation [2, 31–33]

To understand the specific features of CL that underpin this specificity, a plethora of research has been carried out to study CL - cyt c interactions and how they compare to other anionic lipids. Such studies find evidence of an interaction between CL and cyt-c that is both strong and specific, resulting in a distinct complex than cannot be achieved with other anionic lipids [33, 39–42]. The binding to CL can occur through a primary binding site “site A” (**Figure 1b**), which consists of four lysine residues that facilitate an electrostatic interaction with the negatively charged CL [28, 43]. Prior studies have shown CL binding to involve multiple CL lipids [44]. The potency in inducing cyt c peroxidase activity varies among different anionic headgroups and is associated with different extents of protein fold destabilization [45]. PG has been studied in some detail, with prior studies reporting that it seems to preserve a more native-like cyt c structure [46], and facilitates a shallower insertion of cyt c in the lipid bilayer than CL [47]. Also a recent study measuring the femtosecond transient absorption of the heme and its nearby Trp residue suggests that PG and CL cause different degrees of unfolding and change in tertiary structure [48].

A challenge in gaining a comprehensive molecular understanding of the PG- or CL-cyt c complex is that structural inspection of the dynamic nanocomplex is difficult via standard methods of structural biology. Previously we have presented a toolkit of solid-state NMR (ssNMR) approaches to address this challenge for cyt c/CL complexes with CL peroxidase activity [49–52]. Our previous results pointed to CL acting as a dynamic regulator of protein mobility, mediated in large part by the dynamic activation of the heme-binding Ω loop D in cyt c. The highest peroxidase activity, induced by polyunsaturated CL, was also associated with the most pronounced protein mobility [52]. Increases in cyt c mobility upon lipid binding have been revealed by many different studies using an array of different experimental techniques (reviewed recently [14]). One feature of the prior literature is that it includes seemingly conflicting experimental findings regarding the degree of structural and motional changes affecting cyt c. For instance, our own prior work emphasized a greater degree of native-like structure in the bound protein [49–52], relative to findings reported by other groups [14, 29, 44, 45, 53–62]. In the Discussion section we will revisit this issue and examine the potential reasons for these differing findings.

Here, we gain a deeper understanding of the activating mechanism and lipid specificity through experiments on distinct cyt c/lipid complexes spanning a range of protein activity and mobility. We evaluate the connection between binding stoichiometry, lipid peroxidase activation and structural features of cyt c in these complexes. Accounting for measured differences in binding stoichiometry between PG and CL, lipidomics shows the oxidation of polyunsaturated CL to be faster than that of PG, pointing to CL specificity. In a comparison of CL- and PG-bound cyt c by ssNMR the predominant difference is seen in parameters that indicate dynamic differences, alongside modest changes in the NMR chemical shifts. Our data show that distinct lipid species differently regulate cyt c loop dynamics, cause coordinated motions among cyt c foldons and loops, thus regulating protein function. Analogous increases in mobility are also observed upon reducing protein:CL stoichiometries to approach cellular conditions.

## RESULTS

### Stoichiometry of the protein-lipid complex

To examine the formation of protein/lipid complexes by PG and CL with horse heart cyt c, for further functional and structural characterization, we measured cyt c binding to liposomes containing PG or CL. Dissolved cyt c was added to premade large unilamellar vesicles (LUVs) containing DOPC along with a fraction of either DOPG (1,2-dioleoyl-sn-glycero-3-phospho-(1’-rac-glycerol)) or TOCL (tetraoleoylcardiolipin) (**Fig. 1c**). Binding to DOPG-containing LUVs shows a similar trend as for TOCL [52, 63], in that the amount of membrane-bound cyt c increases until a saturation point is reached (**Figure 2a**). Binding saturates when approximately 12 PG molecules are available on the LUV outer layer, facing cyt c. This compares to approximately 6 lipids constituting the saturation point for CL [52]. The lipid area of CL is roughly twice the size of PG [36, 37], and the CL headgroup contains two phosphate groups with similar ionization behavior, implying twice the negative charge of PG (see also **Fig. 1c**) [38]. Thus, both CL and PG feature saturating stoichiometries at approximately twelve negative charges, in a nanodomain with a similar surface area, which in turn matches the footprint of (folded) cyt c [52]. These findings support the common notion that a peripheral interaction between cyt c and anionic lipids can be seen as dominated by electrostatic forces [43, 64].

**Figure 2.**
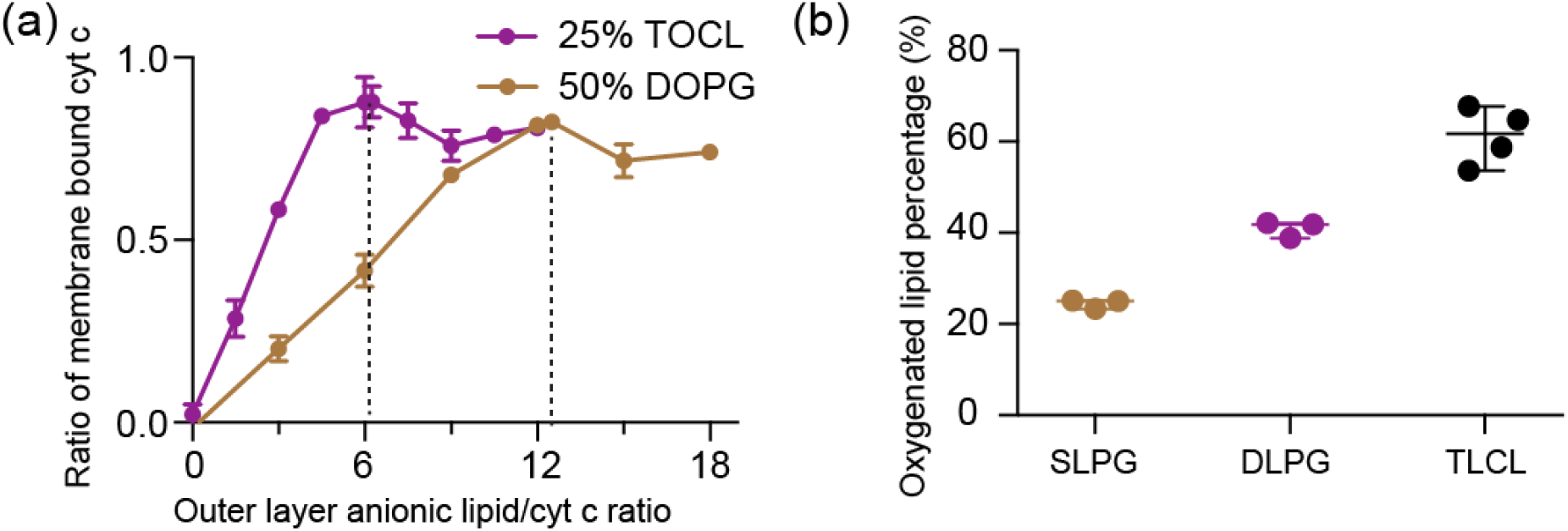
Binding and peroxidase activation by PG and CL containing lipid vesicles. (a) Fraction of membrane-bound cyt c as a function of outer layer anionic lipid/cyt c ratio (“effective” anionic lipid/cyt c ratio). The molar fraction of PG or CL in the DOPG/DOPC and TOCL/DOPC vesicles is indicated. Saturation of cyt c binding to DOPG requires approximately twice the lipids as for TOCL. (b) Comparison of fractional oxygenation of polyunsaturated PG and CL by cyt c/H_2_O_2_ quantified by MS lipidomics. The molar ratio of PG to cyt c was kept at 25:1 and CL to cyt c at 12.5:1 so that the total phosphate to cyt c ratio was the same. The effective anionic lipid/cyt c ratio was 12.5 for PGs and 6.3 for TOCL. The concentration of cyt c was kept at 1 μM, and H_2_O_2_ at 50 μM. Data are presented with mean value and SD.

### PG is a less potent cyt c peroxidase substrate than CL

We further tested whether the formed cyt c - PG complexes attain significant lipid peroxidase activity and compared their activities to the native CL-based complex. Using mass spectrometry lipidomics we measured and quantified cyt c peroxidase activity upon PG binding by the direct detection of oxygenated products of polyunsaturated fatty acid (PUFA) PG: SLPG and DLPG (1-stearoyl-2-linoleoyl-*sn*-glycero-3-[phosphor-rac-(1-glycerol)] and 1,2-dilinoleoyl-*sn*-glycero-3-[Phospho-rac-(1-glycerol)], respectively). Upon incubation with cyt c and H_2_O_2_, oxygenated products of SLPG and DLPG with 1 to 4 oxygens added were observed, with species with two added oxygens dominating, similar to CL oxidation [52]. The oxygenation level of DLPG almost doubles that of SLPG due to the doubled number of unsaturated acyl chains available for oxidation in DLPG. The same number of unsaturated double bonds exist in samples containing DLPG and TLCL, implying a similar chemical reactivity in the peroxidation reaction. Yet, the oxygenation level of DLPG is noticeably lower than that of CL shown in **Fig. 2b**. Thus, PG can act as a substrate but is less potent in inducing cyt c peroxidase activity than CL. We also evaluated cyt c peroxidase activity via the commonly used fluorescence assay of fluorescent product generated by the oxidation of the substrate amplex red. Even when we replaced CL in our LUVs by twice the molar amount of PG (i.e. thereby matching the CL charge and size), we found the peroxidase activity of PG-bound cyt c to be greatly reduced compared to the CL-bound form (**Figure S1a**). Thus, PG mimics the activity of CL, and acts as a potential peroxidation substrate, but is significantly less potent that its CL counterpart.

### CL binding maintains a similar cyt c structure

To better understand the molecular differences in how cyt c behaves in its complex with either PG or CL, we set out to characterize the structure and dynamics of membrane-bound cyt c with ssNMR. To do so, we aim to compare to our prior studies on uniformly ^13^C,^15^N-labeled protein bound to CL-containing membranes [50, 52], and employ analogous methods developed in those studies. An important tool in probing the protein dynamics, and facilitating the effective use of dynamics-sensitive spectral editing (DYSE) techniques [65], was to perform the ssNMR analysis across variable temperatures. This includes measurements at reduced temperatures to suppress the extensive mobility experienced by the protein bound to mono- and polyunsaturated lipid membranes. Even with substantial experimental optimization, the fully labeled lipid-bound protein showed a lot of peak overlap that complicates data analysis. Accordingly, we here performed residue-specific labeling of glycine, isoleucine and leucine residues (^13^C, ^15^N-G, I, L) to reduce peak overlap and improve spectral resolution [52, 66]. These labeled residues were selected for their presence in key segments of the protein, where they act as site-specific probes for secondary and tertiary structure analysis [66]. The selectively labeled proteins were bound to both PG- and CL-containing LUVs, to allow a direct comparison. Cross-polarization (CP) based homonuclear ^13^C-^13^C DARR and heteronuclear ^15^N-^13^C_α_ correlation spectra of PG-bound cyt c are overlaid onto those of CL-bound cyt c in **Figure 3a** and **3b**. 2D ^13^C-^13^C spectra correlate backbone and sidechain carbons of the same residue, and 2D ^15^N-^13^C_α_ spectra show amide nitrogen and C_α_ correlations of the protein backbone. The resonances of the selectively labeled residues match those previously assigned based on studies of the uniformly labeled protein [52]. Directly comparing the PG- and CL-complexes, the spectra are almost identical in peak positions, which is directly indicative of a very similar protein fold in both cases.

**Figure 3.**
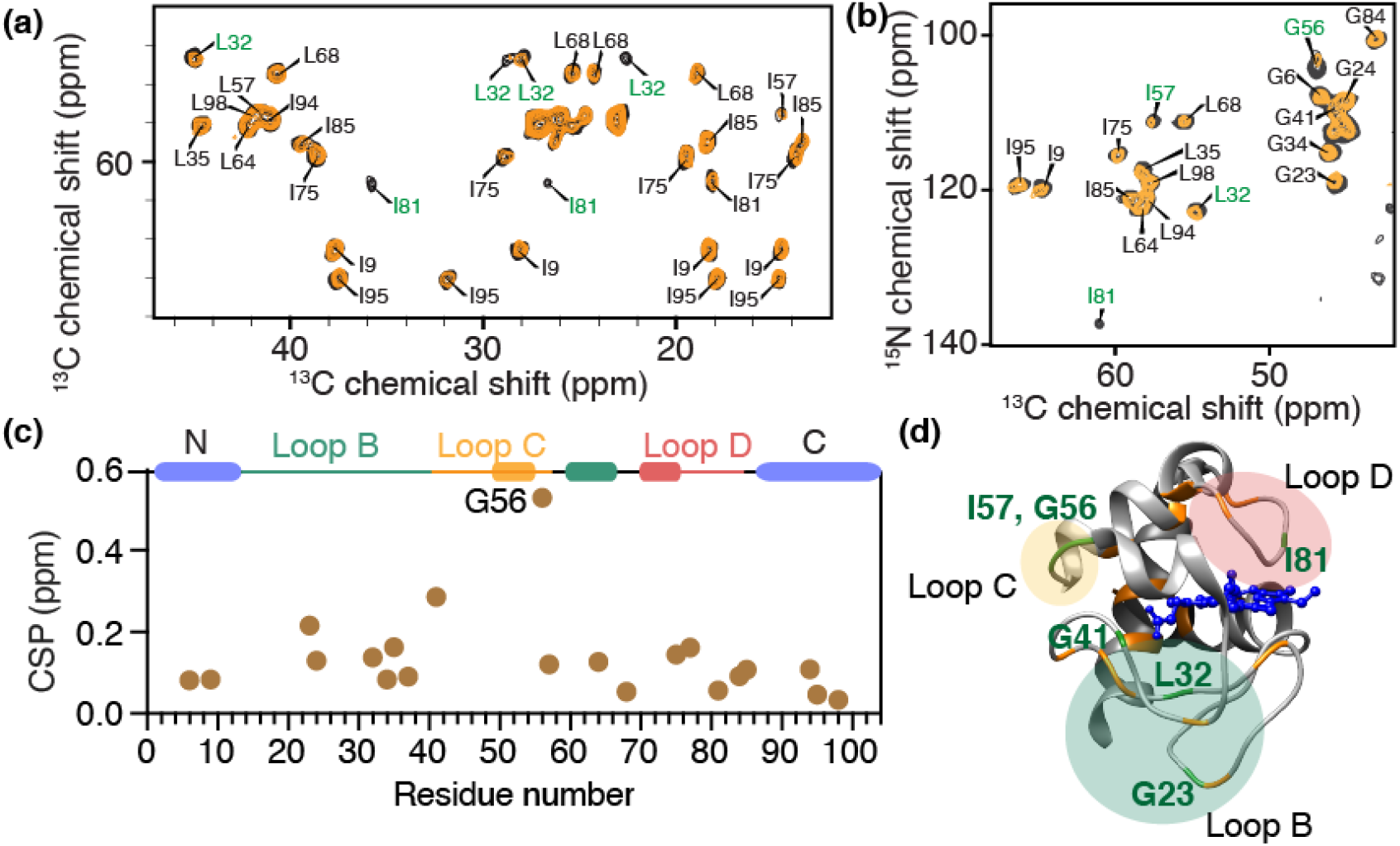
Structural similarities and dynamic perturbations in PG- and CL-bound cyt c. (a) Region from a 2D ^13^C-^13^C CP-DARR ssNMR spectrum showing correlations of backbone Ca with sidechain carbons, and (b) 2D ^15^N-^13^C ssNMR spectrum, for GIL-labeled cyt c bound to DOPC LUVs containing 25% TOCL (yellow) or 50% DOPG (black). GIL-labeled cyt c is selectively labeled with ^13^C, ^15^N- Gly, Ile, and Leu. The total L/P molar ratio is 50 for both, with effective molar PG:cyt c and CL:cyt c ratios of 12.5 and 6.25, respectively. Residues showing prominent peak intensity perturbations are labeled in green. (c) Chemical shift perturbations for GIL-labeled cyt c between the DOPG and TOCL-bound protein, sensing differences between the two lipid complexes. G56 of loop C shows significant perturbation. Helices and foldon units are indicated atop the chart, with colors as in Fig. 1. (d) Horse heart cyt c structure [35], with perturbation sites labeled in green, while orange highlights indicate NMR-observed residues without significant perturbations. The perturbed residues belong to loops B, C and D.

To better understand the structural features of lipid-bound cyt c, we first compare the ssNMR chemical shifts to the shifts of natively folded cyt c studied by solution NMR in reverse micelles [67] (BMRB entry ID 25640) (**Figure S2**). The calculated chemical shift perturbations (CSPs) for PG-bound cyt c are generally below 0.5 ppm, suggesting a high overall similarity between the PG-bound and natively folded state. Residues L68 and I81 show significant CSPs, which is likely due to their proximity to the membrane binding site as also seen for binding to CL-containing membranes [52]. The reader is reminded that potential CSPs in residues that were not labeled would not be detected in the current work, but we also note that the Lys residues constituting binding site A are not resolved, even when labeled. To directly compare the CL and PG bound state of cyt c, CSP analysis *between* those two states is shown in **Figure 3c**. These data reinforce the conclusion that the chemical shifts for cyt c bound to PG and CL are similar throughout all the labeled sites. Glycine residues (G23, G41, and especially G56) show the largest chemical shift changes upon binding to different lipids, perhaps consistent with the small size and innate flexibility of this residue type. These changes point to localized structural differences that can be implicated in recognizing and distinguishing the lipid species. Aside from changes in peak position, we observe intensity differences for residues I81 and L32 in the 2D CP-DARR spectrum and some residues in the 2D NCA spectrum. Here, it is useful to note that in DYSE ssNMR [65] these types of CP-based experiments give optimal signal for the most rigid residues. Then, the lower intensity seen for the CL-bound cyt c shows that cyt c is more dynamic when interacting with CL than PG, with these dynamic differences being residue specific. Residues showing chemical shift and/or intensity perturbations are labeled in **Figure 3d**. The most prominent perturbations are clustered in the loop regions. The perturbed I81 is noteworthy due to its proximity to M80 that coordinates to the heme iron in native cyt-c. Prior studies have shown the M80 residue is to dislodged from its heme coordinating role upon CL binding, as the heme iron goes from hexa- to penta-coordinated [14, 29]. This phenomenon points at a certain degree of destabilization of the native tertiary structure of cyt c.

A recurring question in studies of lipid-bound cyt c relates to the extent to which the native fold is preserved. As one measure of this, we also probed for native-like long-range contacts between α-helices involved in the most stable foldon of cyt c. To do so, we employed 2D ssNMR experiments with long (800 ms) proton-driven spin diffusion (PDSD) ^13^C-^13^C mixing, which allows the detection of inter-residue distances up to 8-9 Å. Local cross-peaks between neighboring residues L94-I95, G84-I85, and between I9 and G6 in the N terminal helix are visible. More interestingly, contacts between G6 and L94 are also observed (**Figure 4a**) consistent with interactions of the N- and C-terminal helices of the protein’s native fold. In its native structure [35] the C_α_ of G6 and C_α_ of L94 are 4.7 Å apart (**Figure 4b**). There are noticeable differences in the intensities of correlations between CL- and PG-bound cyt c. The CL-bound cyt c shows weaker cross peaks, which can result from a more open structure or from the protein being more dynamic, or a combination of structural and dynamic differences. The efficiency of signal transfer in these ssNMR experiments depends on dipolar couplings that are averaged (i.e. reduced) by residual motion [65]. As such, based on these data alone, we cannot conclude a specific distance change. Nonetheless, the detection of inter-residue cross peaks shows that the N- and C-terminal helices are in close contact in both proteolipid complexes, consistent with their role as the most stable foldons of cyt c.

**Figure 4.**
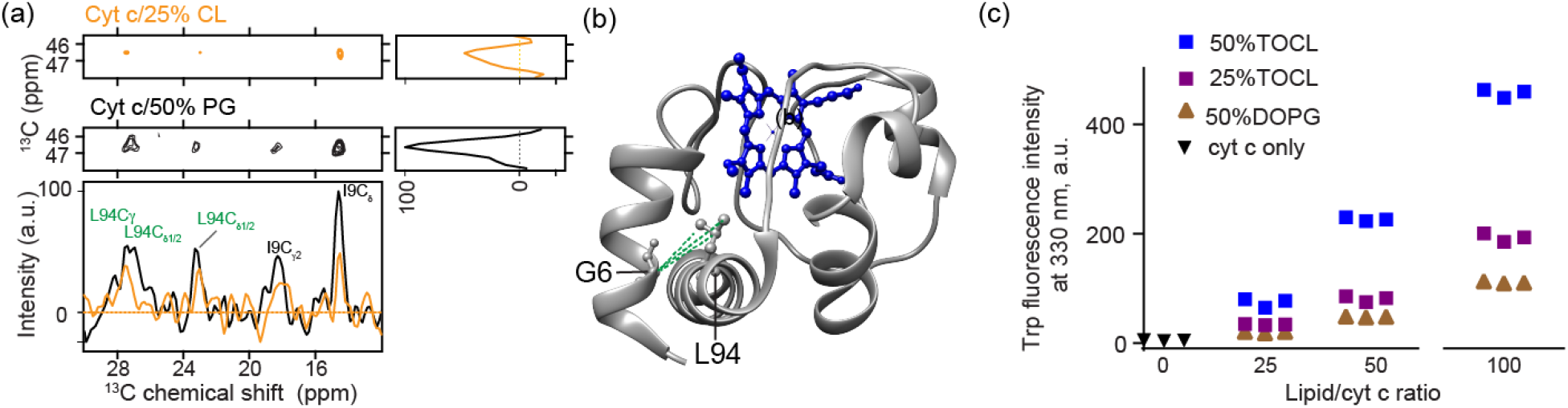
Probing the tertiary fold of cyt c bound to CL and PG LUVs. (a) Extracted regions in 2D ^13^C-^13^C PDSD ssNMR spectra (mixing time 800 ms) showing interhelical contacts between G6 at the N terminus and L94 at the C terminus. The spectrum of cyt c bound to (1:3) TOCL/DOPC liposomes at a LP of 50 is shown in orange and that bound with (1:1) DOPG/DOPC at the same L/P ratio shown in black. 1D slices at the position of G6 Cα showing inter-residual correlations with L94 and I9. 2D spectra are acquired at 265 K, MAS of 10 kHz, and 17.6 T. (b) The N- and C- terminus contacts between G6 and L94 detected are mapped on the cyt c structure in green dashed lines (PDB ID: 1hrc; [35]). (c) CL-induced dynamic changes of cyt c detected by fluorescence spectroscopy. The dynamic changes induced by membranes increase in the order of 50% DOPG, 25% TOCL, and 50% TOCL, and a higher L/P ratio induces more dynamic changes than the lower. Three replicates are presented in the plot. Error bars are within the data point symbols.

### CL is a more efficient dynamic regulator of cyt c than PG

Despite the noted indications of similar structural features upon PG and CL binding, we noticed peak intensity differences in various cyt c residues (**Figures 3a, 3b**, and **Figure 4a**). Such intensity changes are consistent with different extents of dynamic perturbations. The most affected residues are from the three major loops B-D of cyt c (**Figure 3d**). As an orthogonal test of such differences in protein dynamics we monitored the local environment of Trp59 (loop C) via its characteristic fluorescence signal. In natively folded cyt c, the indole ring of Trp59 is close to the heme, which quenches the fluorescence generated upon excitation (**Figure S1b**). When cyt c binds to PG and CL, dynamics affecting the protein increase the Trp-heme distance, resulting in the detection of increased Trp59 fluorescence [29, 44, 56, 68]. **Figure S1b** shows fluorescence spectra of cyt c bound to LUVs with DOPG and TOCL (L/P ratio = 100). The increase of fluorescence intensity can be attributed to a decrease of energy transfer from the Trp to the heme due to an increase of the averaged Trp-heme distance [69, 70]. Compared to PG, a charge-equivalent amount of CL leads to a higher fluorescence emission intensity, showing the protein to be more dynamic. We also observe a slight red shift of the emission, which suggests more water exposure, with the extent of red shift most pronounced for CL. In the context of CL binding, both the extent of red shift and the fluorescence emission intensity correlate to the amount of excess CL (**Figure S1c**). These findings resonate with prior studies that have probed cyt c dynamics and unfolding based on Trp fluorescence, inferring an increased heme-Trp distance with increasing CL:protein ratios [48, 71]. Thus, multiple techniques consistently report increased dynamics in the membrane-bound protein, away from its stable native fold, with the extent of dynamic modulation dictated both by the identity of the anionic lipid substrate *and* the stoichiometry of the protein and lipid.

### Increased CL/cyt c ratios partially destabilize loops but helical structures persist

The lipid stoichiometry is of interest given its reported impact on peroxidase activity and structure of membrane-bound cyt c [14, 50, 52, 55, 72, 73]. We therefore performed ssNMR studies comparing CL-bound cyt c close to the saturating stoichiometry and in presence of an excess of CL. A comparison of 2D ^13^C-^13^C CP-DARR correlation spectra of the CL/cyt c complex at high and low L/P ratios is shown in **Figure 5a** and **5c**. At a high CL/cyt c ratio, we observe more pronounced dynamic perturbations in the membrane-bound cyt c causing many peaks to be missing or perturbed. Interestingly, not all peaks are affected in the same way, since specific peaks are preserved. Supporting the conclusion that these spectral differences stem from dynamic effects, we see that lowering the measurement temperature yielded a spectrum almost identical to that at the lower L/P ratios (**Figure 5b**). One remaining difference is a broader linewidth at the higher L/P ratio, consistent with the low temperature measurement trapping an ensemble with an increased conformational heterogeneity.

**Figure 5.**
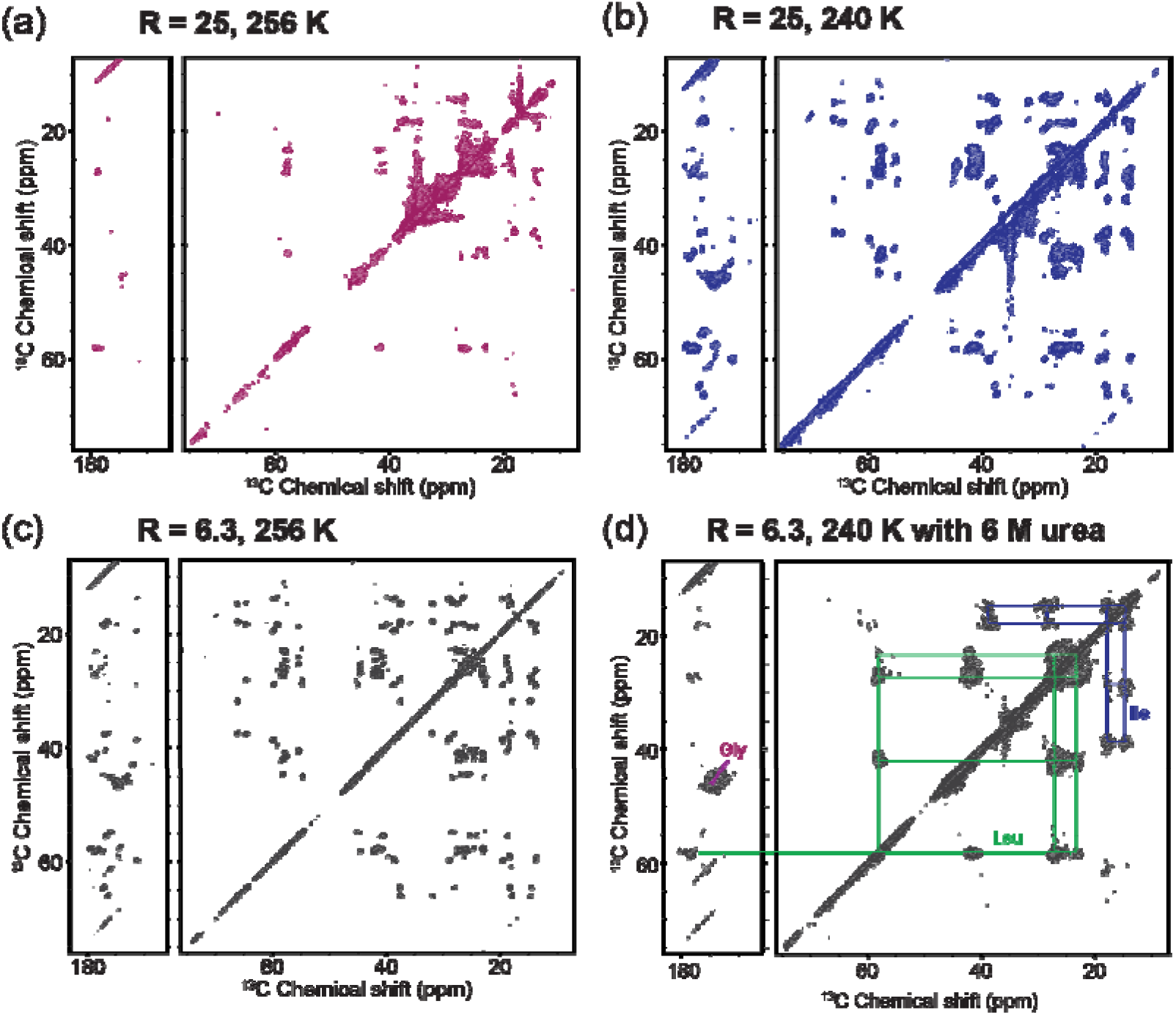
Increased local dynamics for membrane-bound cyt c at higher L/P ratio. 2D ^13^C-^13^C CP-DARR ssNMR spectrum of GIL-labeled cyt c bound to (1:1) TOCL/DOPC vesicles: (a) at an effective CL/cyt c ratio (R) of 25 measured at 256 K; (b) measured at 240K; (c) and at a CL/cyt c ratio of 6.3 at 256 K. The greater excess of CL leads to loss of signals in part of the protein, compared to near-saturating conditions. Suppressing the underlying dynamics at lower temperature (b) leads to recovery of the missing peaks. (d) Chemical denaturation with 6 M urea of membrane-bound cyt c causes the observed signals to feature a narrow chemical shift dispersion and much broader linewidths, consistent with an unfolded state. Urea sample studied at 240 K using (1:1) TOCL/DOPC liposomes at a L/P ratio of 25. All spectra were obtained at 750 MHz (^1^H) and 10 kHz MAS.

Returning to 265K, we can classify residues as being preserved, perturbed, or missing in 2D CP-DARR spectra, in comparison to the analogous data for cyt c at saturated binding. An overlay of the spectra at high and low CL/cyt c ratios is shown in **Figure 6a**. While residue L32 (loop B) is missing, residues G56 and I57 (highlighted in yellow; loop C) are conserved but severely perturbed. Helical residues I9, L64, I95, and I98 are consistently conserved and show minor to slight CSPs, indicating at those sites minor structural changes between the two L/P ratios. Peak intensities reveal dynamic perturbations that increase in the order of the N, 60’s and C helixes being preserved, the C and D loops being perturbed, and B loop being missing (**Figure 6b**). Underpinning the conclusion that these spectral changes are dynamic effects, we saw that they can be largely reverted by modulating the temperature (**Figure 5b**).

**Figure 6.**
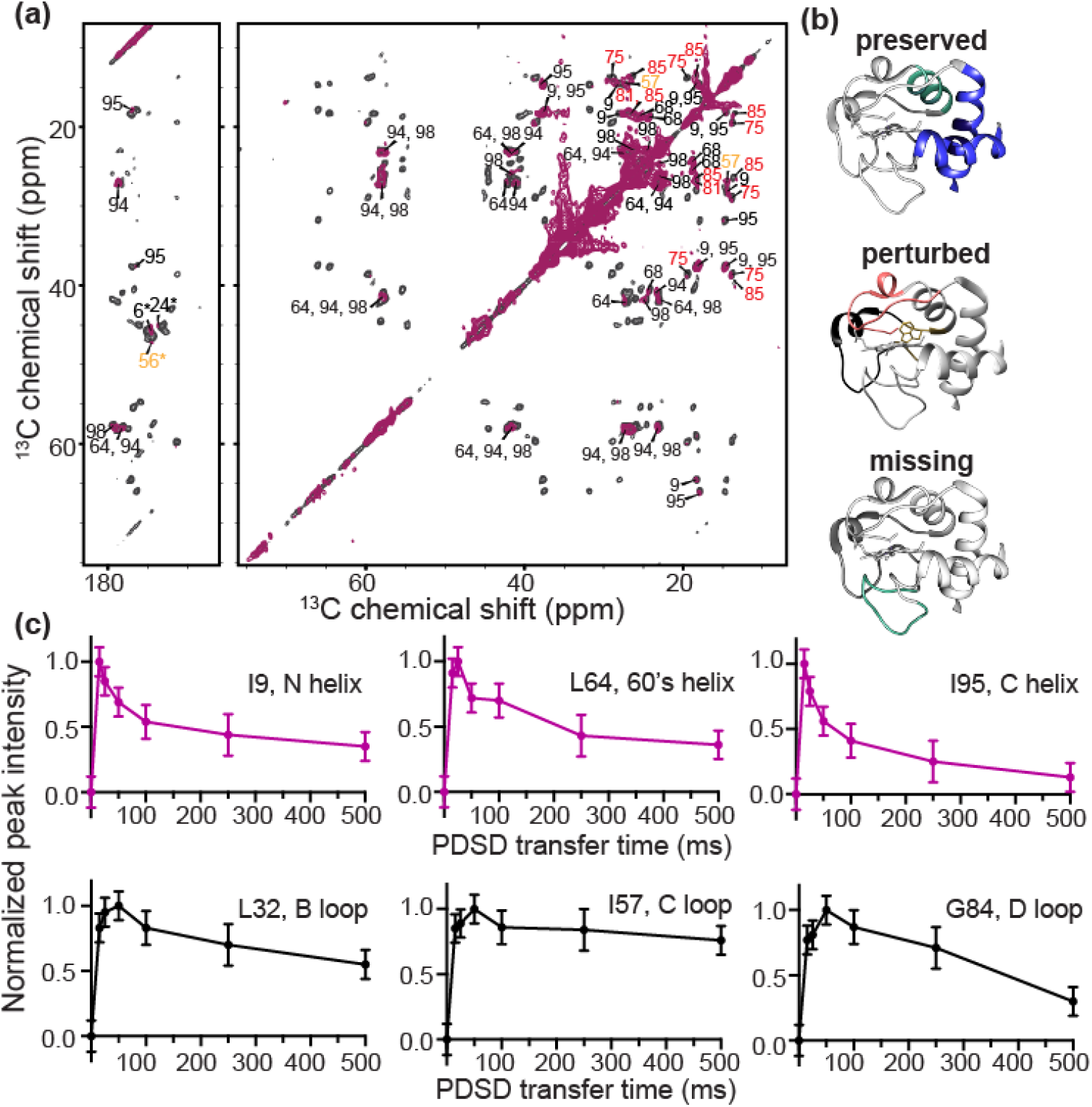
CL binding impacting cyt c foldon dynamics. (a) 2D ^13^C-^13^C ssNMR spectrum of GIL-labeled cyt c bound to (1:1) TOCL:DOPC liposomes at a L/P ratio of 100 (maroon) overlaid with that of cyt c bound to (1:3) TOCL:DOPC liposomes at L/P= 50 LP (black). The effective CL/P ratio R is 25 and 6.3, respectively. Both spectra are acquired at 256 K, 10 kHz, and a magnetic field of 15.6 T. Select residues lack signals in the high CL/P spectrum; the remaining residues are labeled. Those showing intensity and chemical shift perturbations are labeled in red and yellow, indicating the red and yellow foldons. (b) The blue foldon (N-, C-helices) and 60’s helix of the green foldon experience only slow motion at a L/P ratio of 100 (top). Heme-binding Ω loop (red) and 57-60 loop (yellow) exhibit intermediate dynamics (middle). Green foldon (20-35 Ω loop) shows faster dynamics rendering their NMR signals invisible (bottom). (c) PDSD transfer build-up curves for one-bond cross-peaks (C_α_-C’) of α-helix and loop residues, for cyt c bound to (1:3) TOCL:DOPC liposomes at a 50 L/P, acquired at 256 K, 10 kHz, and a magnetic field of 15.6 T. Residues I9, L64, and I95 of helical regions are plotted in purple, and loop residues L32, I57, and G84 in black. The loop residues show increased mobility compared to those in the helices, based on differences in these curves (see text).

In studies of cyt c folding in solution, Englander et al previously noted that destabilization of the foldons by denaturants merely enhanced dynamics present in the native state [10]. We used ssNMR to examine whether some of the pronounced dynamics at high CL/protein ratios are also detectable at near-saturation stoichiometries. To do so, we collected intensity build-up curves for PDSD experiments with mixing times from 0 to 500 ms, for the 50:1 lipid/protein ratio (approximately 6.3 CL per cyt c). Backbone PDSD build-up curves of six residues from different dynamic regions of the protein are shown in **Figure 6c** (with additional data in **Figure S3**). Features such as the maximum intensity, the time of maximum transfer and the rate of intensity decay are affected by motion, through changes in the apparent dipolar coupling strength, relaxation, and chemical exchange effects. The PDSD buildup curves therefore enable a qualitative analysis of protein dynamics, which previously allowed us to distinguish rigid and mobile domains in protein fibrils [74, 75]. I9, L64, and I95 reach maximum transfer at short mixing times and show faster intensity decay in comparison to residues L32, I57, and G84. These features reflect a higher rigidity of the former residues relative to the latter. Although a lack of spectral quality prohibited an analogous analysis for the high L/P sample, these dynamic patterns coincide remarkably with the more pronounced residue-specific dynamic differences apparent in the analysis of peak intensities as discussed above for the excess CL conditions.

### Comparison to complete chemical denaturation

The impact of membranes on cyt c has often been discussed as a process of (partial) unfolding [27, 48, 55, 63], with potential analogy to denaturation of cyt c in solution [71, 76, 77]. In apparent support of this conclusion, an effective CL/cyt c ratio of 25 (L/P=100) causes a similar fluorescence emission to that of GdHCl denatured cyt c (**Figure S1c**). However, the emission wavelength is very different, showing that the extent of water exposure in membrane-bound cyt c remains very limited compared to the fully denatured state. Thus, while it is clear that the normally stable overall fold of cyt c is destabilized upon membrane binding, the phenomenon is distinct from GdHCl denaturation. MAS ssNMR now offers a chance to compare the typical membrane-destabilized state to the effect of chemical denaturation. To do so, we acquired 2D ssNMR spectra of isotope-labeled cyt-c bound to LUVs in presence of 6 M urea (**Figure 5d**). 6M urea has been shown to cause local and global structural changes in cyt c [77]. A prior study reports that the urea should not disrupt the membrane integrity itself [78], a finding supported by dynamic light scattering (DLS) on LUVs in presence and absence of urea (**Figure S4**). The urea-denatured membrane-bound cyt c is highly dynamic, allowing (at 283K) the detection of mobile residues via DYSE experiments based on scalar recoupling methods, e.g. INEPT-TOBSY ^13^C-^13^C spectra that only show flexible residues (**Figure S5**) [65, 79]. In secondary structure analysis these residues lack stable secondary structure. Note that analogous TOBSY spectra of CL-bound cyt c in absence of denaturant lack peaks (data not shown; ref. [50, 52]). Cooling the sample with membrane-bound denatured cyt c suppressed the mobility and allows the acquisition of a 2D CP-DARR spectrum that can be compared to that of non-denatured cyt c. Unlike the non-denatured protein (**Figure 5c**), urea denaturation yields spectra with broad peaks, reflecting highly heterogeneous conformations (**Figure 5d**). As expected for an unfolded protein [80], the spectrum lacks resolution, with peaks being broad and clustered, and the cross-peaks could only be assigned to residue types without site-specific resolution. These data provide a clear visualization of the qualitatively different effect of brute-force denaturation and the more controlled effect of CL-mediated lipid peroxidase activation.

## DISCUSSION

### Differences in lipid peroxidase activation in PG- and CL-cyt c complexes

To better understand the nature of CL-cyt c interactions in pro-apoptotic lipid peroxidase activation we measured how CL and PG mediate membrane binding and lipid peroxidase activation of cyt c *in vitro*. Whilst PG could substitute for CL in facilitating membrane binding, we observed important functional and biophysical differences. Compared to CL, the binding stoichiometry saturated at approximately twice the number of PG lipids per cyt c: a binding capacity of 6.3 CL compared to 12.5 PG. Given the size differences of the lipids, clustering of these lipids would result in the formation of similarly sized nanodomains, which match the footprint of cyt c in its folded state [52]. PG binding triggered cyt c peroxidase activity in amplex red assays and caused PG oxidation as detected by mass spectrometry lipidomics (**Figure 2b**). However, the PG/cyt c nanocomplex was less active in catalyzing peroxidation processes than the CL/protein complex: the efficiency of one equivalent of CL in inducing cyt c peroxidase activity was higher than that of two PG, accounting for their size- and charge differences (**Figure 2b** and **S1a**). Interestingly, the extent of peroxidase activation did not correlate to the extent of binding, showing that structural or motional differences in the bound protein must explain the functional differences. This finding is reminiscent of the impact of different CL ratios on the functional properties of bound cyt c being unexplained by the extent of binding alone [14, 50, 52, 81] as further examined in the current work.

We previously used ssNMR to study the peripherally bound state of cyt c bound to CL-containing membranes [52]. Applying similar methods here, we observed that cyt c bound to PG and CL containing liposomes yields very similar spectra, in terms of peak positions and spectral quality, at least under conditions of comparable charge-corrected P/L ratios (**Figure 2**). The ssNMR peak positions show that loop and helix structures are conserved upon lipid binding. Moreover, we reported here for the first time on structural ssNMR measurements that demonstrate specific inter-helical contacts between the N- and C- termini that form part of the normal tertiary fold (**Figure 4a**). Comparing the PG- and CL-bound spectra, minor CSPs were observed for various residues. These included CSPs for loop residue G56 close to the lipid binding site, and for L32, which is distal to the binding site. Allosteric effects upon binding will be further discussed in later section. The similarity of the ssNMR signals and similar binding behavior (e.g. in terms of the binding saturation at an analogous charge ratio) suggest a qualitatively similar binding mode for CL and PG. Which means that cyt c binds to the ionic lipids retaining significant aspects of its native fold, by interacting with the headgroups through its so-called site A (**Figure 1b**) and additional Lys residues, as reported by us and others [43, 52, 55, 64]. However, even though the two anionic lipids facilitate at-first-glance-similar electrostatic interactions, they nonetheless did not induce the same level of peroxidase activity.

### Distinct dynamics of the PG- and CL-cyt c proteolipid complexes

Our obtained data argue that these functional differences stem from the distinct ways in which the two lipids regulate the dynamics of the protein-lipid complex. In various experimental data we observed that PG binding resulted in a less dynamic protein compared to equivalent CL binding, which itself regulates activity by modulating specific regions of cyt c [52]. The difference in protein dynamics is apparent in our new ssNMR data, and also in Trp fluorescence quenching analysis. This correlation between peroxidase activity and mobility of the PG-cyt c complex is consistent with our prior analysis of how different CL species act as dynamic regulators of cyt c‘s lipid peroxidase activity.

The most pronounced evidence of dynamic differences between PG- and CL-bound cyt c was seen only in particular parts of the protein sequence. These ssNMR-enabled site-specific insights shed more light on the dynamic features of this peroxidase-activate state of cyt c. The enhanced mobility was not affecting the whole protein alike, but rather focused on various key loops. Interestingly, comparing the PG- and CL-bound states we found that their differences were not limited to the heme binding loop D, whose dynamics are crucial for cyt c peroxidase activity. We also noted increased dynamics of loop B and C, representing an apparent allosteric effect of membrane binding, which involves residues on the other side of the protein (**Figure 1a-b**). Worth noting, loop C also contains residue Trp59 which has served as an important probe for the detection of cyt c folding and unfolding based on its heme-proximity based fluorescence quenching [29, 44]. The increased fluorescence and red shift of Trp59 in cyt c can be explained by the increased dynamics upon membrane binding, which cause Trp59 to undergo fast exchange between buried and more exposed states. A similar increase of Trp emission with an increase of anionic charges of the liposomes was recently reported to associate with increased conformational dynamics resulting from partial cyt c unfolding [48]. Consistent with our conclusions of differential dynamics we observed distinct effects on the Trp signal upon PG and CL binding, with the former being much less effective at inducing protein mobility. The mobilizing effect of CL binding was also shown to be strongly dependent on the CL/cyt c ratio, which also impacts on the peroxidase activity. Thus, combined these findings supported the idea that lipid-regulated increases in the plasticity and flexibility of the protein fold are involved in peroxidase activation.

### Membrane binding induces coordinated dynamics in cyt c loops and foldons

We obtained new insights into the nature of these functionally relevant protein dynamics, levering the residue-specific insights enabled by combining 2D ssNMR and targeted isotope labeling. Comparing PG and CL binding and the regulatory effect of the CL/protein ratio yielded complementary insights into the dynamic processes underpinning peroxidase activation. A key finding was that various loops of cyt c are especially implicated in the increased mobility that accompanies the destabilization of cyt c’s native fold upon membrane binding. The helical segments retain a greater degree of rigidity. In this context it is useful to discuss the known folding units, or foldons, or cyt-c (**Figure 1a-b**) [10–13]. We followed the nomenclature introduced by Englander and co-workers, in which the 3D fold of cyt c is formed by cooperative folding of five foldons, which fold in order: blue (N-/C-helices), green (60’s helix and Ω loop B), yellow (a short two-stranded ◻ sheet), red (Ω loop D) and gray (Ω loop C). The stabilizing interaction of N- and C-helixes assists the formation of the first partially folded structural intermediate in the folding pathway of cyt c, forming the blue foldon unit. Next, the blue foldon assists the formation of neighboring green foldons through stabilizing interactions. The subsequent structural intermediates are formed by the adding of foldons in an order of yellow, red and gray. The Ω loop 71-85 (loop D), Ω loop 40-47 (loop C), and loop 20-35 (loop B) assemble in the final stages of folding. Conversely, the destabilization of these loops occurs prior to the unfolding of N- and C-helixes. Englander and co-workers identified a range of conformational intermediates corresponding to the partial folding of foldons. One of the key findings was that these foldons experience H/D exchange (i.e. transient destabilization) even under native conditions, with denaturing conditions merely enhancing these innate instabilities. In the ensemble at native conditions, the high-energy conformational intermediates are present, but exist at low levels compared to the predominantly native structure, as also seen in other proteins [82].

Our results showed that specific cyt c loops are most affected by lipid binding, manifesting in the ssNMR as dynamics-induced changes in intensity. The weakening or disappearance of signals from especially the red and yellow foldons gave a strong indication that the binding of CL favors a partly disassembled and highly dynamic state of cyt c. It is worth noting that loop B is also highly dynamic when binding to the CL-containing membrane, even though it is one of the most stable (i.e. early) foldons under native solution conditions. Thus, the membrane-induced fold destabilization seems to trigger a modified version of the native foldon pathway, although a more detailed understanding of how the lipids modify the (un)folding process requires further experimental and *in silico* study.

An important aspect of our current study is that we observed how an excess of CL lipids leads to more pronounced effects on the protein. A challenge in applying ssNMR to peripheral membrane proteins, like cyt c, is the relatively low sensitivity of the technique [66], especially in comparison to fluorescence or electron spin resonance spectroscopy. This practical challenge implies a trade-off between in-depth characterization and studying low(er) protein concentrations. In our prior ssNMR studies we targeted a regime where cyt c was close to saturating the CL binding sites, representing a higher L/P ratio than feasible in ssNMR studies in the 1990s. Here we observe, both in Trp fluorescence assays and ssNMR spectra a clear impact of a greater excess of CL on the ssNMR spectra and dynamics of the bound protein. We observed a loss of many signals from the late-stage foldons, even while signals from the most stable foldons are preserved. These preserved foldons constitute almost all of the α-helical secondary structure in the protein. This is fully consistent with prior FTIR studies by ourselves and others that show the preservation of secondary structure in membrane-bound cyt c [50, 57, 83, 84]. We observed that the dynamic changes at higher CL excess reflect an increase in pre-existing mobility of particular foldons of the membrane-bound protein, even at the more stable CL-saturating conditions (**Fig. 6**).

The protein dynamics on the membrane showed a hierarchical order with the N, 60’s and C helixes being most stable (**Fig. 7a**), in apparent analogy to their roles as early foldons. One exception is the destabilization of loop B by membrane binding, which contrasts to its usual concerted folding with the 60’s helix. The heme-binding D loop, 50’s helix and its adjacent C loop show increased dynamics on the NMR timescale. These findings support the importance of D loop dynamics as implicated in peroxidase activation, which is perhaps not surprising given that this loop harbors the important Met80 heme ligand [52]. These insights deepen our understanding of the reported impacts of the protein:lipid ratio on protein mobility and peroxidase activity [14, 50, 52, 55, 72, 73], revealing changes in the hierarchical dynamics that affect the membrane-bound protein.

**Figure 7.**
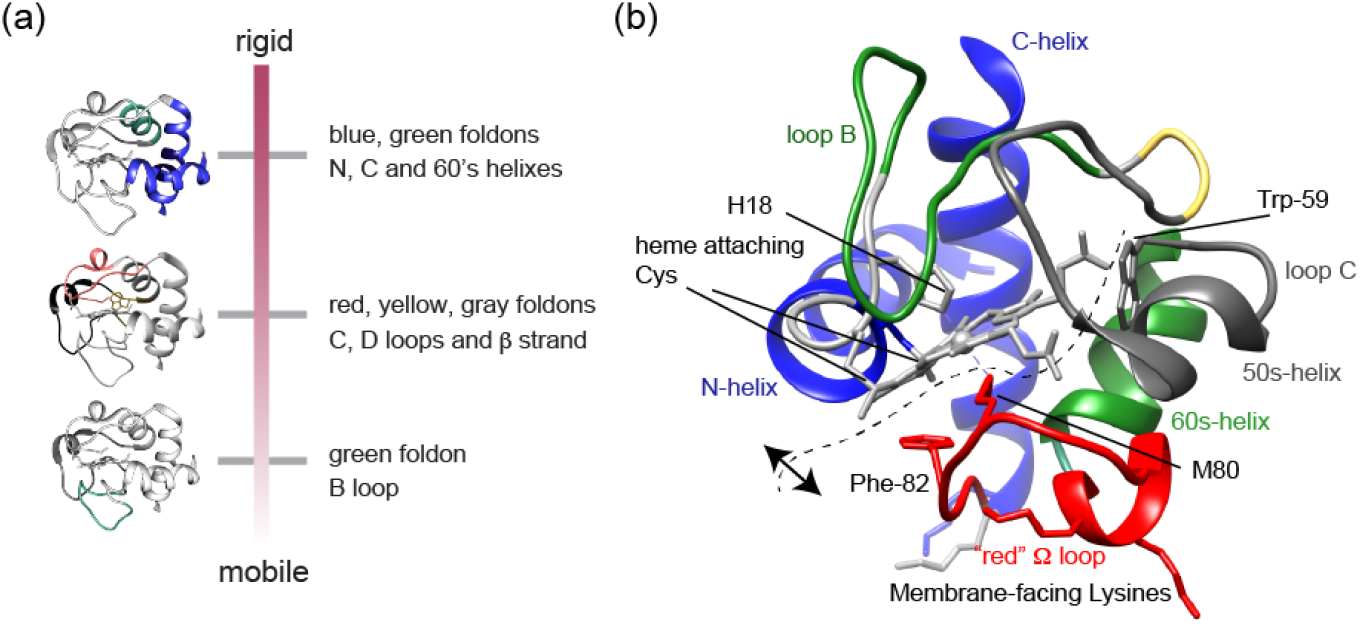
Hierarchical dynamics of folding units of cyt c upon binding to CL-containing membranes. (a) Overview of hierarchical dynamics detected by ssNMR. (b) Molecular structure of cyt c highlighting key residues and segments. The dashed line indicates the hypothesized opening of the CL-bound protein fold, such that it would preserve the stable blue foldons with associated heme (top/left), while increasing the spacing between heme and residues Met80, Phe82 and Trp59 (right/bottom), as detected spectroscopically. The range of motion depends on the experimental conditions (e.g. lipid composition).

### The dynamic ensemble of membrane-bound cyt c

In our initial studies of cyt c binding to CL-containing membranes we noted the similarities of NMR spectra for bound and unbound cyt c, which argued against a large scale unfolding under the conditions studied [50]. This result contrasted with the findings from other studies and techniques that emphasized an unfolding of the protein on the membrane. The current experiments finetune our understanding of the protein fold in the membrane-bound state. We see a pattern where certain lipids (e.g. PG), lower temperatures and high membrane surface occupancy favor a native-like fold that is not undergoing much internal motion. This is seen in ssNMR spectra but also in Trp fluorescence data. On the other hand, more disruption from this native-like state was observed as we moved to CL lipids, lower protein-to-lipid ratios, and ambient temperatures. These conditions yielded ssNMR spectra where many peaks were missing in CP-based experiments that detect parts of the protein that are more ordered and rigid. In dipolar-based ssNMR experiments we probed protein dynamics on the micro- to millisecond timescales, which are of critical importance for membrane protein function [85, 86]. The ssNMR results provide valuable insights into the differential motions of cyt c foldons upon interaction with CL, within this timescale regime. We observed a hierarchical pattern in the disappearance of peaks, indicating differential dynamics in cyt c foldons (**Figure 7a**). It is instructive to relate our data to those previously reported from other techniques. Spectroscopic studies of the heme and residues internal to cyt c (e.g. Trp59 and Phe82; **Figure 7b**) show that the tertiary fold of the protein becomes more open and flexible [43, 56, 61, 71]. I.e. the distance between the heme and Trp59 or Phe82 increases. Moreover, experiments have shown the displacement of the native heme ligands upon membrane binding [61, 71, 87]. At the same time, various data (including FTIR and ssNMR) show that α-helical structures are preserved [50, 52, 83, 84]. Here we observed that the N- and C-terminal helices were retained and interacted. Notably, this constitutes the most stable foldon that also features heme interactions through the covalent Cys bonds and His18 (**Figure 7b**). In contrast, the interactions with other parts of the protein were becoming less stable, and subject to dynamic exchange processes. Thus, while there are native-like states in the membrane-bound ensemble, these native states are in exchange with more open structural states [29, 45, 53, 58, 59, 61, 62, 71]. Even for the more open states in the membrane-bound ensemble, it would appear that key features (i.e. secondary structures, helix-heme interactions and certain foldon elements) are preserved, while other parts are more disturbed (e.g. loop structures facing the membrane). This fits with spectroscopic observations and explains repetitive observations of reversibility in the membrane-induced destabilization [61, 62, 64, 88, 89]. We propose that this perspective unifies much of the available data. It is important to also note that cyt c-lipid interactions vary greatly depending on the experimental conditions, such as the pH, ionic strength and other parameters that we have not explored in the current report.

### On lipid-induced allosteric mobility affecting cyt c loop C

As noted above, the ssNMR data identify increased mobility not only in membrane-facing regions, but also in other segments such as loop C. Thus, it is possible that loop C dynamics play a role in the activation of cyt c’s peroxidase function. This kind of role is supported by recent studies of cyt c mutants showing that mutations in loop C can also regulate cyt c peroxidase activity [90–92]. Lei et al. determined the X-ray crystal structure of the A51V variant of cyt c, which shows only minor changes in tertiary structure compared to the WT protein, but exhibits several folds of increase in peroxidase activity [90]. A G41T mutation causes increased mobility of both loops C and D, resulting in loop D destabilization [92]. Worrall and coworkers have also shown that increased dynamics of loop C in a several mutants increase the population of a peroxidase-active state, in which Met80 of loop D is disassociated [54, 91]. Although these studies were carried out on cyt c in the absence of lipids, they highlight the important role of loop C in modulating the peroxidase activity implicated in apoptosis. We see also in the membrane-bound state increases in loop C and D dynamics as a result of CL binding, which may be one of the mechanisms by which membrane binding regulates the protein function. Low frequency motion is a known feature of biological membranes, causing membrane-associated proteins to be subjected to similar kinds of motion [93, 94]. The kinds of protein dynamics on the microsecond to millisecond timescales or longer, which are implicated by our ssNMR data, have been associated with important biological functions, including ligand binding and molecular recognition, enzymatic activity, and protein folding [95].

It is well known that membrane protein dynamics are important for protein functions. So far, ssNMR studies of the interplay between membrane biophysics and membrane protein structure and dynamics are largely limited to transmembrane proteins [85]. In most cases, protein dynamics invoke the interconversion between the ground state and functional state of the transmembrane domain. Even though the conformational dynamics of peripheral membrane binding protein cyt c have been studied by several spectroscopic methods, these methods probe either a single site or the protein as one entity. Our ssNMR studies provide site-specific structure and dynamics of cyt c upon its binding with ionic lipids. We demonstrated how peripheral membrane binding can regulate protein functions through the modulation of foldon dynamics. Higher peroxidase activity is correlated to higher dynamics, e.g. upon binding to CL rather than PG, or as a function of the lipid:protein stochiometry. Our findings substantiate a clear distinction between complete unfolding of the protein (which also facilitates peroxidase activity) and the coordinated dynamics of the lipid-cyt c complex that catalyzes lipid peroxidation. The interplay of this peripherally bound membrane protein with its bound lipids controls the characteristics of its functional state, reflecting a dynamic regulation mechanism that is sensitive to intricate details of the lipid species and the membrane biophysical features.

## MATERIALS AND METHODS

### Materials

Common chemicals were purchased from Fisher Scientific (Pittsburgh, PA) or Sigma-Aldrich (St. Louis, MO). Horse heart cytochrome c (natural abundance) was purchased from Sigma-Aldrich (catalog number C7752). Phospholipid stocks were obtained from Avanti Polar Lipids (Alabaster, Alabama), including dioleoyl phosphatidylcholine (1,2-dioleoyl-*sn*-glycero-3-phosphocholine, DOPC; C18:1), tetraoleoyl cardiolipin (1’,3’-bis[1,2-dioleoyl-*sn*-glycero-3-phospho]-*sn*-glycerol (sodium salt), TOCL; C18:1), 1-stearoyl-2-linoleoyl-*sn*-glycero-3-[phospho-rac-(1-glycerol)] (sodium salt), (SLPG: 18:0-18:2 PG), 1,2-dilinoleoyl-*sn*-glycero-3-[phospho-rac-(1-glycerol)] sodium salt (DLPG: 18:2 PG), and 1,2-dioleoyl-*sn*-glycero-3-phospho-(1’-rac-glycerol) sodium salt (DOPG: 18:1 PG). 1’,3’-bis[1,2-dimyristoyl-*sn*-glycero-3-phospho]-glycerol, sodium salt, (TMCL: 14:0 cardiolipin), 1-palmitoyl(D31)-2-oleoyl-*sn*-glycero-3-phosphocholine and 1-palmitoyl(D31)-2-oleoyl-*sn*-glycero-3-[phospho-rac(1-glycerol)] were used as internal MS standards.

### Liposome preparation

Mixed lipid vesicles of DOPG with DOPC were prepared as previously described [50, 52]. DOPG and DOPC in chloroform were mixed in a 1:1 molar ratio and dried under N_2_ gas for 15 – 20 min and placed under vacuum in a desiccator overnight to remove residual solvent. The dried lipid film was resuspended in HEPES buffer (20 mM HEPES, pH 7.4) to form multilamellar vesicles by vortexing. Afterwards, the liposome suspension was flash frozen by insertion into a cold bath made from dry ice and ethanol, and thawed completely by transferring to a hot water bath at approximately 52 °C. The freeze-and-thaw cycle was repeated 10 times. The final DOPG/DOPC (1:1) liposome stock was obtained by extrusion through a 200 nm filter unit (T&T Scientific Corp, Knoxville, TN) for 11 times. The typical total lipid concentration was 10 mg/mL. Other liposomes with different molar ratios used throughout our studies, including SLPG/DOPG/DOPC (0.33:0.67:1), DLPG/DOPG/DOPC (0.33:0.67:1), TOCL/DOPC (1:1), and TOCL/DOPC (1:4) liposomes, were prepared consistently by the same protocol unless stated otherwise.

### Dynamic Light Scattering (DLS) analysis

DLS measurements were obtained on a Zetasizer Nano instrument (Malvern Panalytical Ltd, UK) in order to test vesicle integrity in presence and absence of urea. A suspension of 100 μM liposomes in HEPES buffer (20 mM HEPES, pH 7.4) was prepared in a disposable sizing cuvette. Urea stock solution in HEPES buffer was added to reach a final concentration of 6 M. An acquisition time of 60 – 80 s was determined by the instrument. Ten repeated measurements were performed for each sample. Representative size distribution plots were generated by the instrument software to show particle size and distribution.

### Measurement of cyt c Binding to Negatively Charged Lipid Vesicles

Liposomes consisting of either an equal molar amount of DOPG and DOPC, or 25 mol-% TOCL and 75% DOPC were prepared following the protocol described above. Cyt c stock solution was prepared by dissolving cyt c crystals in sample buffer (20 mM HEPES, 100 μM diethylenetriaminepentaacetic acid (DTPA), pH 7.4) Different ratios of LUVs and cyt c (with the protein concentration kept at 5 μM) were incubated in the sample buffer at room temperature for 15 min in ultracentrifugal tubes (Beckman Coulter, Indianapolis, IN). The effective PG/cyt c molar ratio (R) ranges from 0 to 48, and the effective CL/cyt c ratio is from 0 to 24. The effective lipid/protein ratio R represents the calculated ratio of lipids on the outer layer of the vesicles relative to the protein. The unbound cyt c was separated from liposome-bound cyt c by ultracentrifugation at 435,000 g for 2 hours at 4 °C in an Optima TLX ultracentrifuge with a TLA-100 rotor (Beckman Coulter). Immediately after the ultracentrifugation, the supernatant containing unbound cyt c was carefully removed and saved for UV-vis spectrophotometer (Beckman Coulter) measurement. Cyt c solutions without adding liposome or centrifugation were prepared in parallel as a reference. UV-vis absorbance at 409 nm was recorded for samples (S) and references (S_0_), and the ratio of membrane-bound cyt c was calculated as 1-S/S_0_ for plotting as a function of the outer layer anionic lipid/cyt c molar ratio R. Experiments were performed in duplicate.

### Tryptophan Fluorescence Measurements

For Trp fluorescence measurements, the vesicles prepared as described above were further sonicated five times for 30 s on ice using an ultrasonic probe tip sonicator (Cole-Palmer Ultrasonic Homogenizer, 20 kHz, Cole-Palmer Instrument Company, Vernon Hills, IL). Cyt c (10 μM) was incubated with liposomes of different compositions and at different ratios. Trp fluorescence was measured using a PCI steady-state photo counting spectrofluorometer (ISS Inc. Champaign, IL) using quartz cuvettes. The excitation wavelength was 290 nm. Each measurement was performed in triplicate.

### Mass-spectrometric Analysis of Polyunsaturated Lipid Oxygenation Catalyzed by Cyt c

We measured and quantified the oxygenation of polyunsaturated lipids catalyzed by cyt c in presence of H_2_O_2_ using a mass spectrometry lipidomics method developed and reported before [52]. Mixed liposomes containing SLPG, DOPG, and DOPC at a ratio of 0.33:0.67:1 and those incorporating DLPG instead of SLPG were prepared following the procedure described above. Liposome stock was added to cyt c stock solution in HEPES buffer (20 mM HEPES, 100 uM DTPA, pH 7.4) to reach a lipid to protein ratio of 50 and a final cyt c concentration of 1 μM. The oxidation was initiated by adding freshly prepared H_2_O_2_ stock to a final concentration of 50 μM, with further additions every 15 min during 1 hour of incubation on a shaker at 25 °C. Control samples which contained lipid only, contained LUVs and cyt c, or contained LUVs and H_2_O_2_ were prepared in parallel. The extent of lipid oxidation and lipid oxidation species were detected and characterized by mass spectroscopy.

LC–MS of lipids. Lipids were extracted by Folch procedure with slight modifications, under nitrogen atmosphere at all steps. LC–ESI–MS/MS analysis of lipids was performed on a Dionex HPLC system (using the Chromeleon software), consisting of a mobile phase pump, a degassing unit and an autosampler (sampler chamber temperature was set at 4 °C). Phospholipids were separated on a reverse phase Acclaim C30 column (3 μm, 150 × 2.1 mm (ThermoFisher Sientific)) at a flow rate of 0.060 mL/min. The column was eluted using gradient of solvent system consisting of mobile phase A (acetonitrile/water/triethylamine, 45/5/2.5 v/v) and B (2-propanol/water/trimethylamine, 45/5/2.5 v/v). Both mobile phases contained 5 mM acetic acid and 0.01% formic acid. The Dionex HPLC system was coupled to a hybrid quadrupole-orbitrap mass spectrometer, Q-Exactive (ThermoFisher Scientific) with the Xcalibur operating system. The instrument was operated in negative ion mode (at a voltage differential of −3.5–5.0 kV, source temperature was maintained at 150°C). Analysis of LC–MS data was performed using the software package Compound Discoverer (ThemoFisher Scientific). The resolution was set up at 140,000 that corresponds to 5 ppm in *m/z* measurement error. *M/Z* values for PLs and their oxidation species are presented to 4 decimal points.

The incubation of liposomes containing partial SLPG or DLPG with cyt c /H_2_O_2_ triggers the oxidation of SLPG and DLPG. Under the conditions used, the production of oxygenated products containing 1 to 4 oxygens is detected. The total amount of oxygenated SLPG and DLPG is estimated as 244 and 410 pmol/nmol, respectively. The oxidation of TLCL under the same condition was previously reported [52] and plotted as a reference in Fig2b.

### Identification of PG hydroxy- and hydroperoxy-molecular species

Identification of oxidized PG molecular species was performed by their mass-charge ratio (MS1) and fragmentation properties (MS2) using ESI-MS analysis. In negative ion mode, PG produces singly anionic charged [M-H]^−^ ion with corresponding signals of non-oxidized SLPG and DLPG with *m/z* 773.5321 and 769.5008, respectively. Trace amount of the sodium acetate adducts of SLPG and DLPG ions with *m/z* 851.5025 and 855.5338, respectively, was also observed. ESI-MS assessment of PG oxidation products reveals the accumulation of different hydroxy-, hydroperoxy-, dihydroxy- and hydroxy/hydroperoxy- derivative of SLPG and DLPG as evidenced by the appearance of signals with *m/z* 789.5287; 785.4974; 805.523; 801.4923; 833.4825; 821.5172 and 817.4880, respectively.

LC-MS^2^ fragmentation analysis of SLPG revealed that species containing 9-OOH-LA (*m/z* 805.523; Rf 21.1 min), 13-OH-LA (*m/z* 789.5287; RF24.5) and 9-OH -LA (*m/z* 789.5287; Rf 25.4 min) were the major SLPG oxidation products. Among those hydroperoxy- species were predominant. Similarly, oxidation of DLPG by cyt C/H_2_O_2_ resulted in the major product containing 9-OOH-LA in one acyl chain (*m/z* 801.4923, Rf 14.7 min). In addition, 9-OH-LA (*m/z* 785.4974; Rf 18.8min) and 13-OH-LA (*m/z* 785.4974; 18.3 min) and 9-10-epoxy-LA (*m/z* 785.4974; Rf 20.7 min) were detected as well. Furthermore, LC-MS analysis revealed the presence of di-hydroxy-species containing OH groups in both acyl chains of DLPG (*m/z* 801.4923; Rf 11.9 min). Hydroxy-hydroperoxy- molecular species were localized in one acyl chain only for both types of PG and were observed in SLPG (*m/z* 821.5172, Rf 20.83) as well DLPG (*m/z* 817.4880; Rf 10.7 min). In addition, di-hydroxy- molecular species in DLPG were found in both acyl chains in C9 position (*m/z* 833.4825; 9.8 min).

### Expression and Purification of ^13^C, ^15^N-G, I, L Specifically Labeled Cyt c

The expression and purification of ^13^C, ^15^N-G, -I, -L labeled horse heart cyt c were carried out similarly to reported procedures [50]. To produce ^13^C, ^15^N-G, I, L specifically labeled cyt c, cells were grown at 24 °C in minimal media. When an optical density of 1.0-1.2 was reached, the media was supplemented with 0.1 g/L of each of the (unlabeled) amino acids except for glycine, isoleucine and leucine, which were added in their ^13^C, ^15^N-enriched form (Sigma) at 0.1 g/L. The media were also supplemented with 0.5 mM FeCl_3_ and 580 μM unlabeled 5-aminolevulinic acid hydrochloride. Protein overexpression was induced with 1 mM IPTG. The cells continued to grow at 24 °C for 4 h before being pelleted down by centrifugation and resuspended in 25 mM sodium phosphate buffer (pH 6.5) with 0.02% sodium azide. The cells were lysed by homogenizer and the cell debris removed by centrifugation. The protein purification was carried out using a cation exchange HiTrap SP column (GE Healthcare, Chicago, IL) with phosphate buffer (25 mM sodium phosphate, 10% (v/v) glycerol, 0.02% sodium azide, pH 6.5) and a 0 – 1.0 M NaCl salt gradient. The column eluate containing cyt c was further purified through a gel filtration Superdex 75 16/600 column (GE Healthcare, Chicago, IL) phosphate buffer (25 mM sodium phosphate, 150 mM NaCl, 5% glycerol, 0.02% sodium azide, pH 6.5). The oxidized form of cyt c was obtained by incubation with 5-fold molar amount of potassium hexacyanoferrate (III) (Sigma-Aldrich) for 15 min. Excess K_3_Fe(CN)_6_ was removed through buffer exchange in 10 kDa Amicon Ultra-15 centrifugal filters with HEPES buffer (20 mM HEPES, pH 7.4).

### SSNMR Sample Preparation

Liposomes containing 50% DOPG and 50% DOPC were prepared in HEPES buffer (20 mM HEPES, pH 7.4) according to the method described above. Next, ^13^C, ^15^N-G, I, L-specifically labeled cyt c stock solution was added to the liposome stock at a molar lipid/protein ratio of 50, and the mixture was vortexed at room temperature for 15 min. Note that this ratio reflects the total lipid content, which is different from the “effective” lipid:protein ratio (R) that was used above to evaluate the accessible lipids on the outer surface of the vesicles. The protein/liposome complexes were pelleted into a Bruker 3.2 mm thin wall rotor using a home-built rotor packing tool at 325 k g for 2 – 5 hours [51]. The sample containing ^13^C, ^15^N-G, I, L-specifically labeled cyt c bound to LUVs containing 25% TOCL and 75% DOPC was prepared at a total L/P ratio of 50. Each sample contained approximately 3 mg of the labeled protein.

For the denaturing studies, LUVs containing 25% TOCL and 75% DOPC were mixed with ^13^C, ^15^N-G, I, L-specifically labeled cyt c at a L/P molar ratio of 50. Urea solution was subsequently added to the abovementioned mixture to reach a final urea concentration of 6M. After incubation at room temperature for 20 min, the urea-denatured cyt c/membrane complexes were packed into a Bruker 3.2 mm thin-wall MAS rotor using the rotor packing tool mentioned above. Upon centrifugation into the MAS rotor, the binding percentage was determined based on UV-VIS analysis of the supernatant, as described above, which revealed a reduced binding percentage of about 70% compared to 95% in the absence of 6 M urea.

### NMR Experiments, Data Processing, and Analysis

NMR measurements were carried out on a Bruker Avance III 750 MHz wide-bore spectrometer and a Bruker Avance 600 MHz wide-bore spectrometer (Bruker, Billerica, MA). A typical ^1^H 90° radio frequency (RF) pulse was 3 μs. The MAS ssNMR samples were spun at 10 kHz. In 2D ^13^C-^13^C Dipolar Assisted Rotational Resonance (DARR) and ^15^N-^13^C (NCA) correlation experiments [96, 97], the magnetization from ^1^H to ^13^C or ^15^N was first transferred through cross polarization (CP). The RF strengths on ^1^H and the heteronucleus were matched to the Hartmann-Hahn condition. A typical contact time of 1 ms and 2 ms was used for ^1^H-^13^C and ^1^H-^15^N CP transfer, respectively. The ^15^N-^13^C contact time was 4 ms for the second CP of the NCA experiment. A DARR spectrum with a mixing time of 25 ms was acquired for chemical shift assignments. To detect long-range interactions, a PDSD spectrum was acquired with a ^13^C-^13^C mixing time of 800 ms. ^1^H decoupling power of 83 kHz was applied during acquisition and evolution periods. The ssNMR signals of immobilized and dynamic parts of the sample were probed via dynamic spectral editing (DYSE) experiments [65]. Flexible signals were identified in 1D J-coupling-based ^13^C spectra acquired using rotor-synchronized refocused insensitive nuclei enhanced polarization transfer (INEPT) ^1^H-^13^C transfers, along with 2D spectra with ^13^C-^13^C transfers enabled by the P9^1^_3_ total through bond correlation spectroscopy (TOBSY) pulse sequence [79], at a MAS rate of 10 kHz. A series of PDSD experiments with mixing times of 0, 15, 25, 50, 100, 250, and 500 ms was acquired. Peak area integrations were normalized and plotted as a function of mixing times, and compared to reference data on rigid samples [75]. Detailed experimental parameters are included in table S1 in the supplemental information.

NMR spectra were processed with the NMRPipe software package [98]. 2D spectra were processed involving apodization with 90° sine-bell function, linear prediction of the indirect dimension with once the original data size, zero filling, and Fourier transformation. The chemical shifts were referenced to dilute aqueous 2,2-dimethylsilapentane-5-sulfonic acid (DSS) and liquid ammonia, using adamantane ^13^C chemical shifts as an external reference, as previously described [50, 75]. NMR data analysis was performed using the Sparky program (T.D. Goddard and D. G. Kneller, Sparky 3, UCSF).

## ABBREVIATIONS

CL: cardiolipin
CP: cross polarization
CSP: chemical shift perturbation
cyt c: cytochrome c
DARR: Dipolar Assisted Rotational Resonance
DLPG: 1,2-dilinoleoyl-*sn*-glycero-3-[Phospho-rac-(1-glycerol)]
DOPC: 1,2-dioleoyl-*sn*-glycero-3-phosphocholine
DOPG: 2-dioleoyl-*sn*-glycero-3-phospho-(1’-racglycerol)
DSS: 2,2-dimethylsilapentane-5-sulfonic acid
DTPA: diethylenetriaminepentaacetic acid
DYSE: dynamic spectral editing
IMS: intermembrane space
INEPT: insensitive nuclei enhanced by polarization transfer
L/P: lipid:protein ratio
LUV: large unilamellar vesicle
MAS: magic angle spinning
MIM: mitochondrial inner membrane
MOMP: membrane permeabilization
MS: mass spectrometry
PDSD: proton-driven spin diffusion
PG: phosphatidylglycerol
RF: radio frequency
ROS: reactive oxygen species
ssNMR: solid-state nuclear magnetic resonance
SLPG: 1-stearoyl-2-linoleoyl-*sn*-glycero-3-[phospho-rac-(1-glycerol)]
TLCL: tetralinoleoyl cardiolipin (1’,3’-bis[1,2-dilinoleoyl-*sn*-glycero-3-phospho]-*sn*-glycerol)]
TOBSY: through bond correlation spectroscopy
TPPM: two-pulse phase-modulated decoupling
TOCL: tetraoleoyl cardiolipin (1’,3’-bis[1,2-dioleoyl-*sn*-glycero-3-phospho]-*sn*-glycerol).

## Acknowledgements

We thank Mike Delk for advice and support with the NMR measurements. The authors acknowledge funding support from the National Institutes of Health R01 GM113908 to P.C.A. v.d. Wel, P01 HL114453 and U19AI068021 to V.E. Kagan, and NIH instrument grant S10 OD012213-01 for the 750 MHz NMR spectrometer.

