## Supporting Information for "Activation of Cytochrome C Peroxidase Function Through Coordinated Foldon Loop Dynamics upon Interaction with Anionic Lipids"

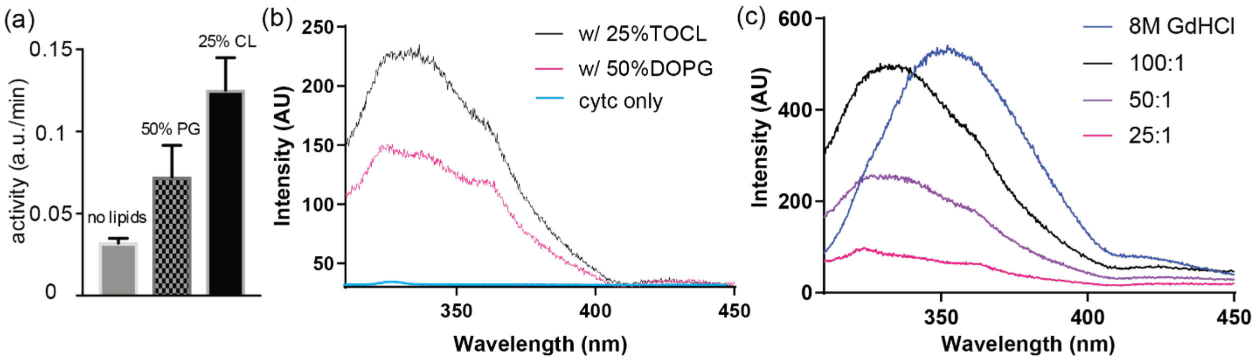


**Figure S1. Membrane-binding induced peroxidase activity c and dynamic change of cyt c by fluorescence-based measurements.** (a) Peroxidase activity of cyt c induced by liposomes containing 50% DOPG/50% DOPC and 20% TOCL/80% DOPC, as measured by in amplex red assay. A total lipid to protein (L/P) molar ratio of 25 was used. (b) Comparison of Trp fluorescence spectra of cyt c-DOPG and cyt c-TOCL complexes at a total L/P ratio of 100:1. The Trp fluorescence of dissolved cyt c without lipids is highly quenched, as also shown. (c) Trp fluorescence spectra of cyt c-TOCL complexes at different L/P ratios (total molar L/P ratio indicated in legend). Mixed liposomes containing 25% TOCL and 75% DOPC were used. The fluorescence intensity increases with an increase of the L/P ratio, indicating a partial dynamic opening of the protein's 3D fold. Fully denatured cyt c in 8 M GdHCl (in absence of lipids) is shown as a reference. Aside from intensity changes, also the frequency of the maximum is changing, indicating different extents of water exposure (see main text).

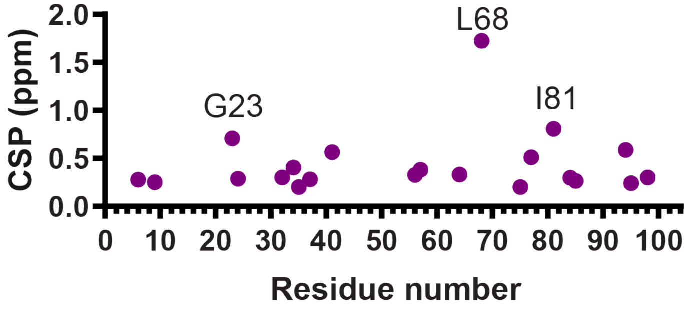


**Figure S2. Chemical shift perturbations of cyt c bound to PG-containing liposomes.** The plotted CSP values are between unbound soluble cyt c (BMRB entry ID 25640; [1]) and cyt c bound to mixed lipid LUVs with a composition of 50% DOPG and 50% DOPC and an effective molar ratio of PG to cyt c of 12.5.

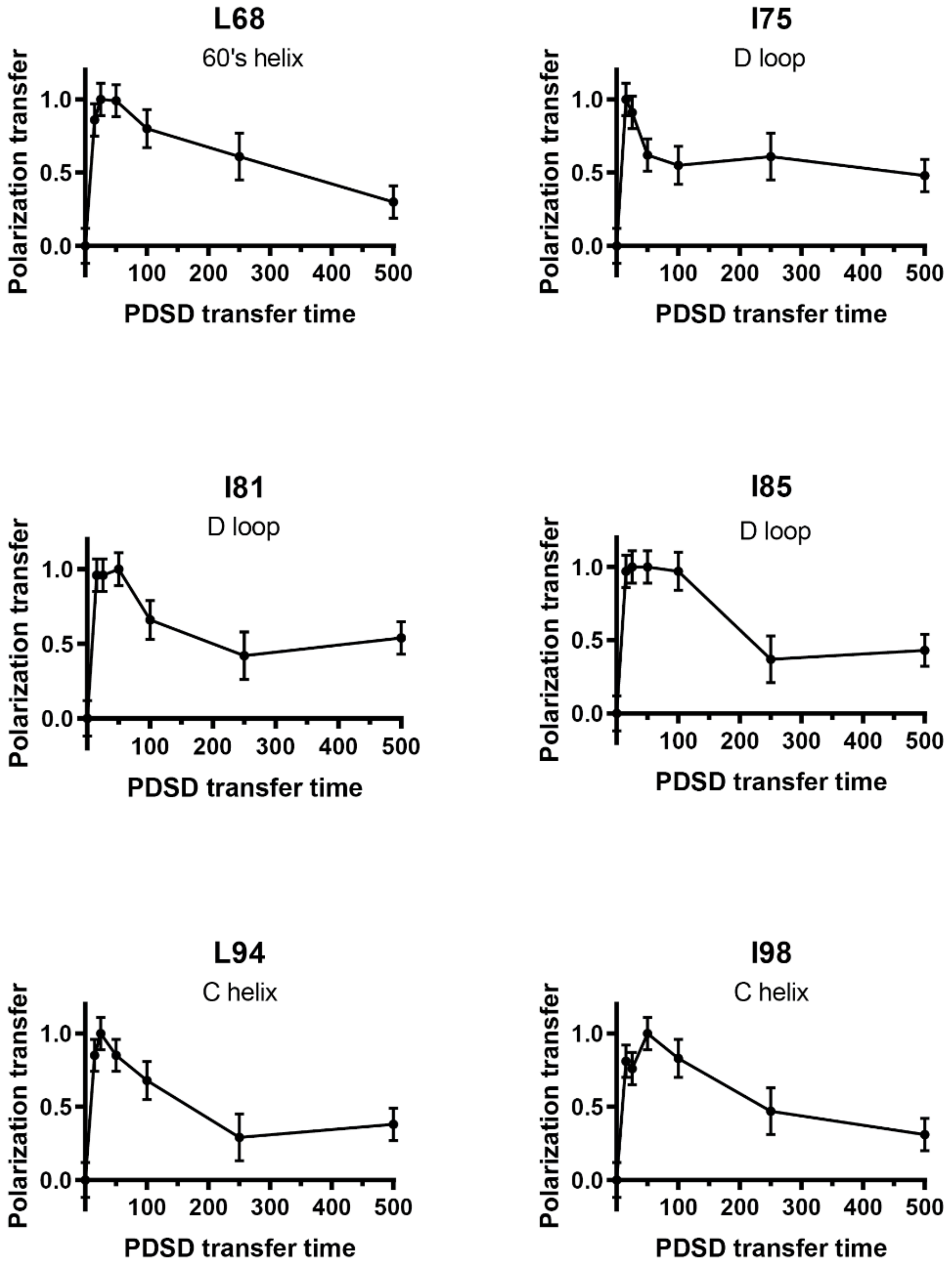


**Figure S3**. **PDSD intensity build-up curves for one-bond cross-peaks of α-helix and loop residues**. These data were obtained from a series of PDSD experiments with different mixing times, applied to labeled cyt c bound to (1:3) TOCL:DOPC liposomes at a 50 L/P. The experiments were performed at 256 K, 10 kHz MAS, and a magnetic field of 15.6 T.

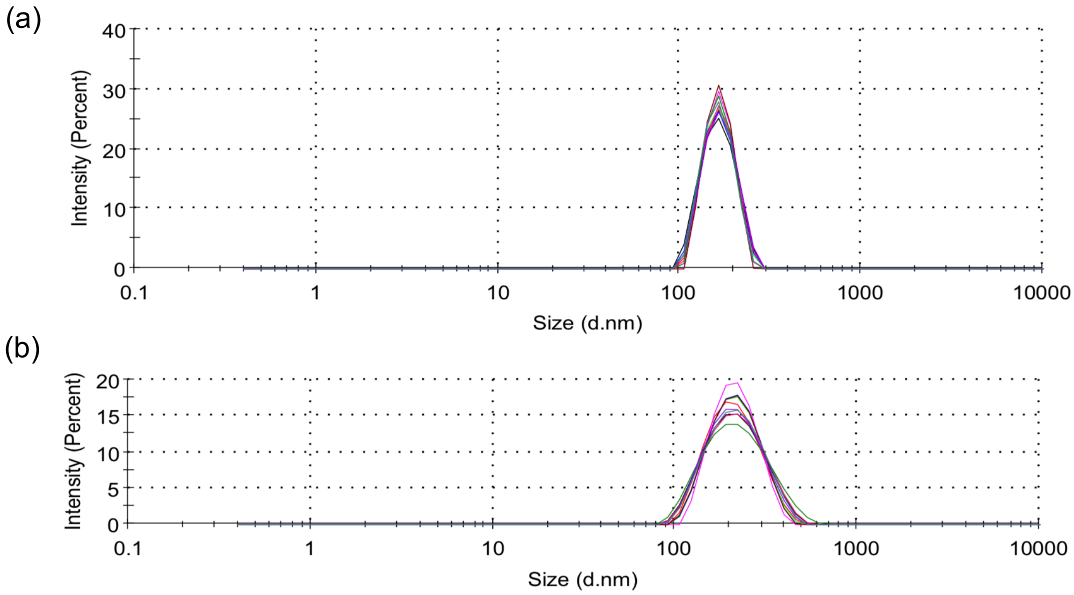


**Figure S4**. **DLS analysis of lipid vesicles with and without added 6M urea**. The integrity of membrane vesicles in presence of 6 M urea was evaluated by particle size measurement from DLS. Particle size and distribution of liposomes containing 50% TOCL and 50 DOPC, (a) without and (b) with 6M urea. Ten repeated DLS measurements of each sample are plotted in the graph. Binary (1:1)TOCL/DOPC liposomes show monomodal distribution with an averaged particle size of about 170 nm. In presence 6M urea, membrane particles are still monodispersed with similar particle size. The distribution is slightly broader than that without urea. However, there is no indication of membrane disruption such as the formation of micelles of much smaller size.

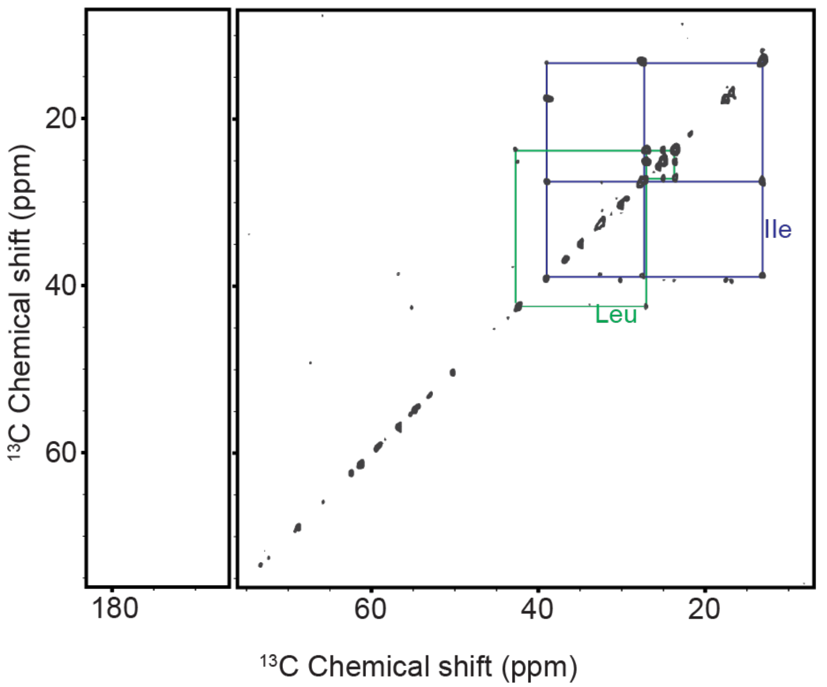


**Figure S5**. **2D INEPT-TOBSY spectrum of chemically denatured cyt c bound to (1:1) TOCL/DOPC LUVs.** This 2D spectrum was acquired with $^{13}C$,$^{15}N$-GIL-labeled cyt c, bound to the LUVs at a CL/cyt c ratio of 25, in the presence of 6 M urea. Cross-peaks from the side chains of Leu and Ile residues are marked. Since such INEPT-TOBSY spectra only detect highly flexible residues with relatively long $T_2$ relaxation times, these data reveal the presence of mobilized and flexible residues in the membrane-bound denatured protein. Corresponding spectra of membrane-bound cyt c in absence of urea lack these cross peaks. The spectrum was acquired on a 750 MHz NMR instrument at a temperature of 283 K and using 10 kHz MAS.

**Table S1. Experimental conditions for MAS ssNMR experiments[1].**

| NMR Sample | Figure | Expt. | NS | Temp | MAS | RD | $t_1$ evol. | Mix. | CP time |
|---|---|---|---|---|---|---|---|---|---|
| | | | | K | kHz | s | ms | ms | ms |
| GIL-labeled cyt c bound to (1:1) DOPG:DOPC at L/P=50 | 3a | 2D DARR | 64 | 265 | 10 | 2 | 6ms (500 x 24.1us) | 25 | 1.7 ms |
| GIL-labeled cyt c bound to (1:3) TOCL:DOPC at L/P=50 | 3a | 2D DARR | 128 | 265 | 10 | 2 | 6ms (500 x 24.1us) | 25 | 1.7 ms |
| GIL-labeled cyt c bound to (1:1) DOPG:DOPC at L/P=50 | 3b | 2D NCA | 640 | 265 | 10 | 2 | 10ms (80 x 250us) | NA | 1.5 ms / 4.2 ms |
| GIL-labeled cyt c bound to (1:3) TOCL:DOPC at L/P=50 | 3b | 2D NCA | 1920 | 265 | 10 | 2 | 10ms (80 x 250us) | NA | 1.5 ms / 4.4 ms |
| GIL-labeled cyt c bound to (1:1) DOPG:DOPC at L/P=50 | 4a | 2D PDSD | 192 | 265 | 10 | 2 | 6ms (500 x 24.1us) | 800 | 1.7ms |
| GIL-labeled cyt c bound to (1:3) TOCL:DOPC at L/P=50 | 4a | 2D PDSD | 192 | 265 | 10 | 2 | 6ms (500 x 24.1us) | 800 | 1.7ms |
| GIL-labeled cyt c bound to (1:1) TOCL:DOPC at L/P=100 | 5a, 6a | 2D DARR | 192 | 256 | 10 | 2 | 3.6ms (300 x 24.1us) | 25 | 1.0 ms |
| GIL-labeled cyt c bound to (1:1) TOCL:DOPC at L/P=100 | 5b | 2D DARR | 512 | 240 | 10 | 2 | 3.6ms (300 x 24.1us) | 25 | 1.0 ms |
| GIL-labeled cyt c bound to (1:3) TOCL:DOPC at L/P=50 | 5c, 5b | 2D DARR | 64 | 256 | 10 | 2 | 6ms (500 x 24.1us) | 25 | 1.7ms |
| GIL-labeled cyt c bound to (1:1) TOCL:DOPC at L/P=25 w/ 6M urea | 5d | 2D DARR | 128 | 240 | 10 | 2 | 3.6ms (300 x 24.1us) | 25 | 1.0 ms |
| GIL-labeled cyt c bound to (1:3) TOCL:DOPC at L/P=50 | 6c | 2D PDSD | 64 | 256 | 10 | 2 | 6ms (500 x 24.1us) | 0 | 1.7 ms |
| GIL-labeled cyt c bound to (1:3) TOCL:DOPC at L/P=50 | 6c | 2D PDSD | 64 | 256 | 10 | 2 | 6ms (500 x 24.1us) | 15 | 1.7 ms |
| GIL-labeled cyt c bound to (1:3) TOCL:DOPC at L/P=50 | 6c | 2D PDSD | 64 | 256 | 10 | 2 | 6ms (500 x 24.1us) | 25 | 1.7 ms |
| GIL-labeled cyt c bound to (1:3) TOCL:DOPC at L/P=50 | 6c | 2D PDSD | 64 | 256 | 10 | 2 | 6ms (500 x 24.1us) | 50 | 1.7 ms |
| GIL-labeled cyt c bound to (1:3) TOCL:DOPC at L/P=50 | 6c | 2D PDSD | 64 | 256 | 10 | 2 | 6ms (500 x 24.1us) | 100 | 1.7 ms |
| GIL-labeled cyt c bound to (1:3) TOCL:DOPC at L/P=50 | 6c | 2D PDSD | 64 | 256 | 10 | 2 | 6ms (500 x 24.1us) | 250 | 1.7 ms |
| GIL-labeled cyt c bound to (1:3) TOCL:DOPC at L/P=50 | 6c | 2D PDSD | 64 | 256 | 10 | 2 | 6ms (500 x 24.1us) | 500 | 1.7 ms |
| GIL-labeled cyt c bound to (1:1) TOCL:DOPC at L/P=25 w/ 6M urea | S3 | 2D INEPT-TOBSY | 160 | 283 | 10 | 2 | 6.6ms (250 x 53us) | 6 | NA |

*[1]) Abbreviations: NS, number of scans per $t_1$ point; MAS, magic angle spinning rate; RD, recycle delay; $t_1$ evol., $t_1$ evolution time = (real and imaginary $t_1$ points) X $t_1$ increment time; Mixing, $^{13}C$-$^{13}C$ or $^{1}H$-$^{1}H$ mixing time (ms); Temp. Temperature; $^{1}H$ decoupling power during evolution and acquisition using the two-pulse phase modulation scheme (TPPM) was 83 kHz for all experiments.*

### ***References cited in the Supplemental Data***


[1] O'Brien ES, Nucci NV, Fuglestad B, Tommos C, Wand AJ. Defining the Apoptotic Trigger: The interaction of cytochrome c and cardiolipin. J Biol Chem. 2015;290:30879-87.
